## Supplementary Information for "ERK2-topoisomerase II regulatory axis is important for gene activation in immediate early genes"

**This PDF contains:**

Supplementary Figures 1–7

Supplementary Tables 1 and 2

**Supplementary Data contains:**

Supplementary Data 1: Mass spectrometry data- deTOP2B control

Supplementary Data 2: Mass spectrometry data- deTOP2B with ERK1

Supplementary Data 3: Mass spectrometry data- deTOP2B with ERK2

Supplementary Data 4: Mass spectrometry data- deTOP2B with ERK1m

Supplementary Data 5: Mass spectrometry data- deTOP2B with ERK2m

Supplementary Data 6: Mass spectrometry data- spectrum

Supplementary Data 7: Cryo-EM validation report

**A**

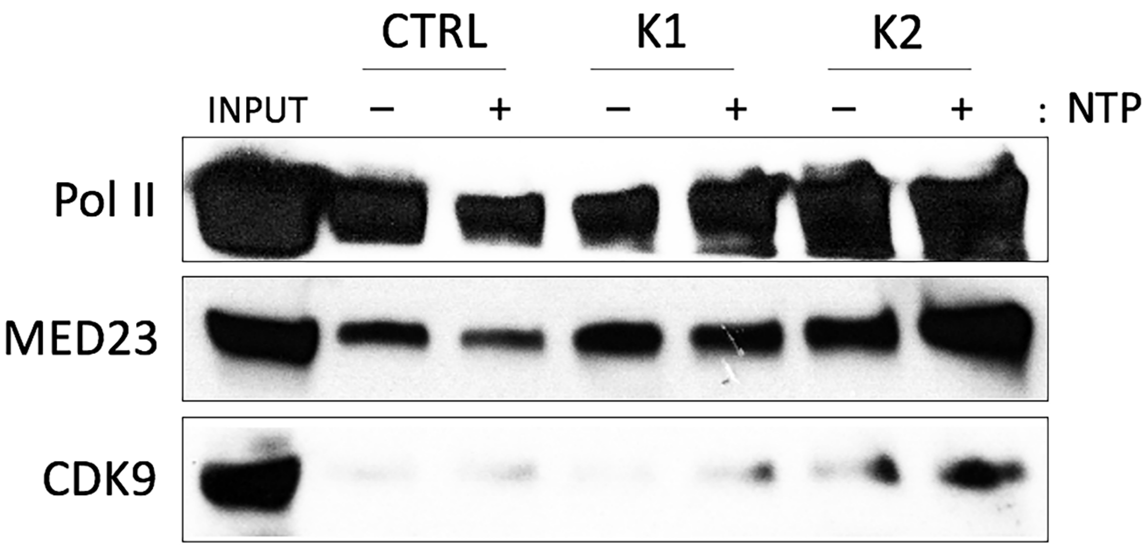

**B**

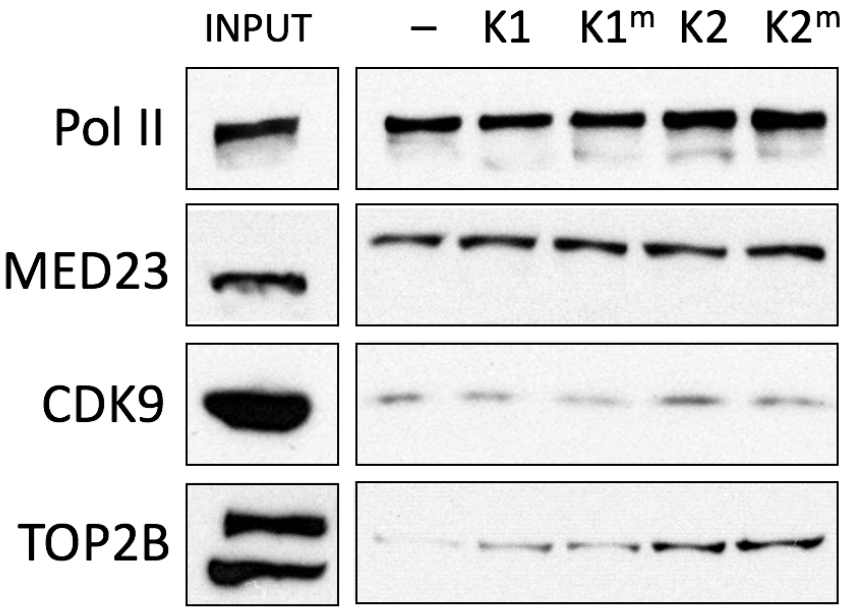
INPUT
–
K1
K1m
K2
K2m
Pol II
MED23
CDK9
TOP2B

**C**

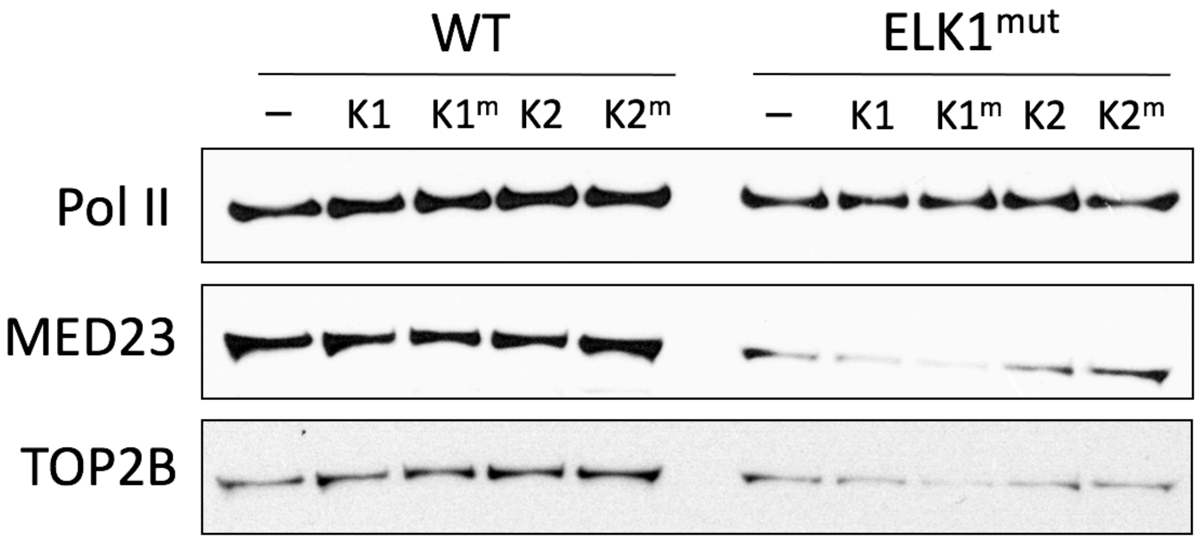
WT
ELK1mut
–
K1
K1m
K2
K2m
Pol II
MED23
TOP2B

**Supplementary Figure 1. ERK2 activates EGR1 transcription.**

(**A**) Immobilized template assay results showing more prominent effects of ERK2 on recruiting Pol II, MED23, and CDK9 to the *EGR1* TSS. ERKs were supplemented with HeLa NE and before NTP addition. CTRL, ERK storage buffer only without ERKs.
(**B**) Immobilized template assay results indicating that ERK2 and ERK2m are more effective to recruit transcription factors, such as Pol II, MED23, TOP2B, and CDK9, on the *EGR1* template than ERK1 and ERK1m (upper panel). Input (NE), 20%. –, ERK storage buffer only control.
(**C**) The stimulating effects by ERK2 and ERK2m is dependent on ELK1 binding on the template. –, ERK storage buffer only control.

**A**

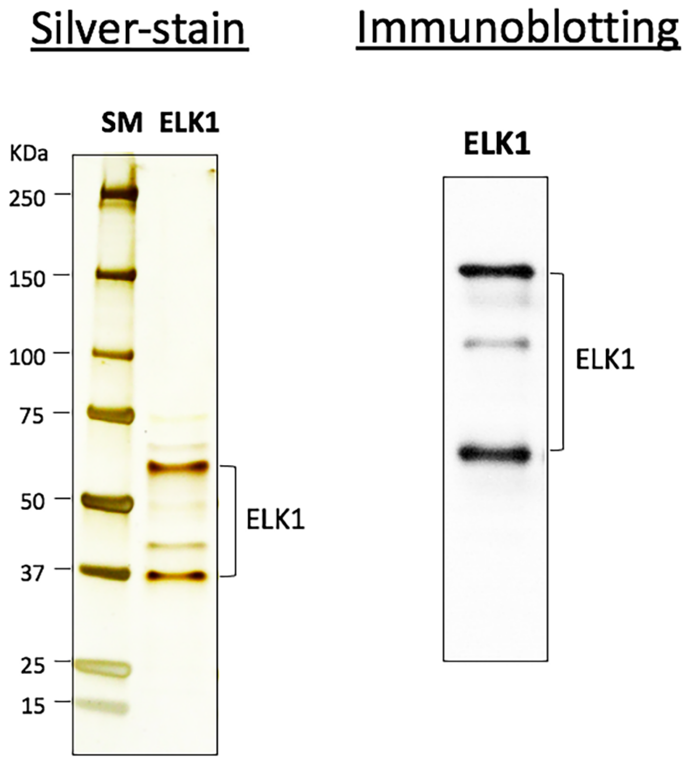
Silver-stain
Immunoblotting
SM
ELK1
KDa
250
150
100
75
50
37
25
15
ELK1
ELK1
ELK1

**B**

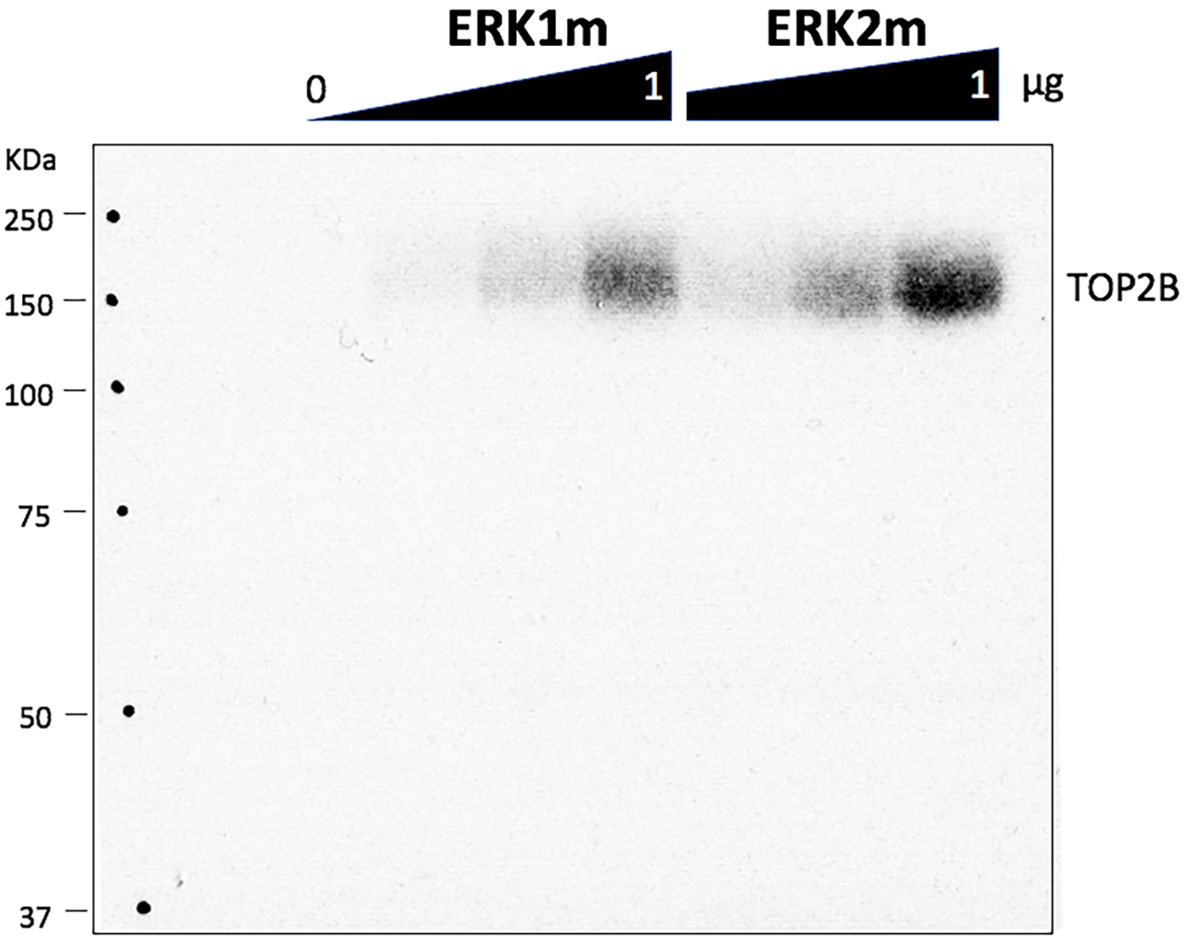


**Supplementary Figure 2. *In vitro* kinase assay and autoradiography of a SDS PAGE gel.**

(**A**) Recombinant ELK1 that was purified from *E. coli* and used for the *in vitro* kinase assays in this study is shown on a SDS-PAGE gel stained with silver (left) and on an immunoblot (right).
(**B**) Titration of ERK1m and ERK2m in the *in vitro* kinase assay showing a dose-dependent increase of phosphorylation of TOP2B. Nine μg of TOP2B and 0, 0.25, 0.5 and 1 μg of ERK1m or ERK2m were incubated for 1 h at 30 °C. A $^{32}P$ autoradiography of SDS-PAGE gel.

**A**

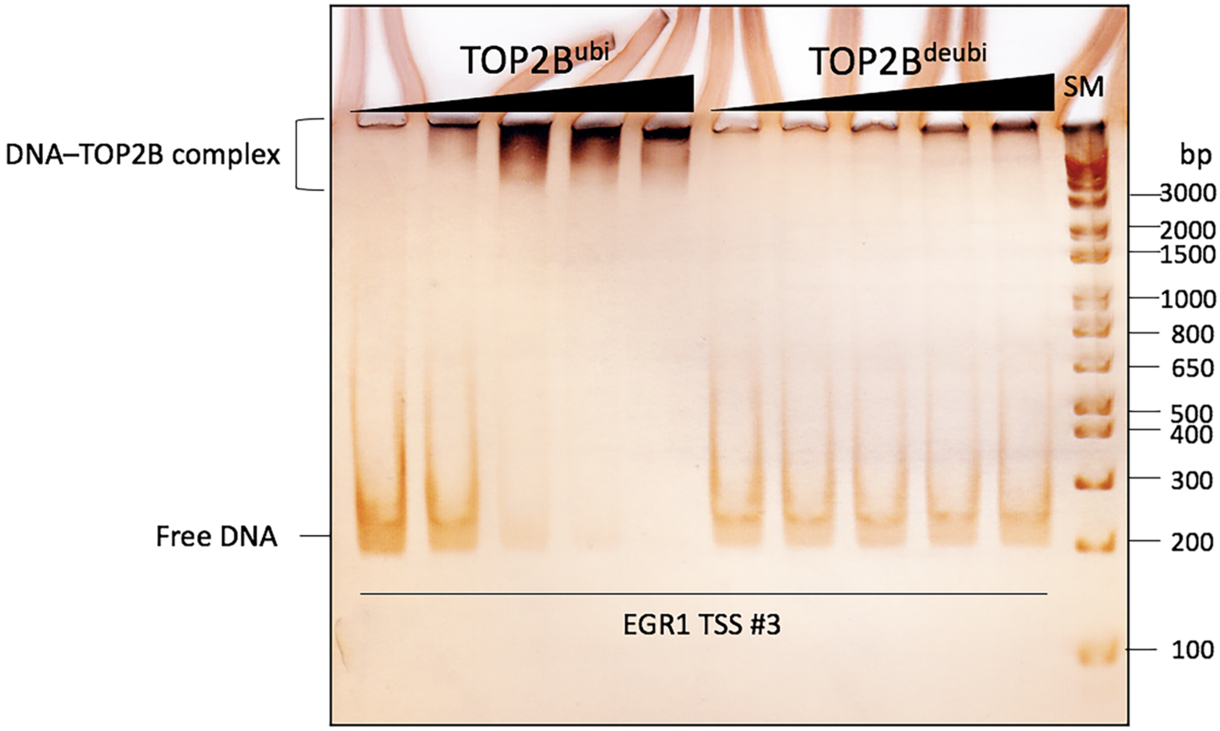
TOP2B^ubi
TOP2B^deubi
SM
DNA–TOP2B complex
bp
3000
2000
1500
1000
800
650
500
400
300
200
100
Free DNA
EGR1 TSS #3

**B**

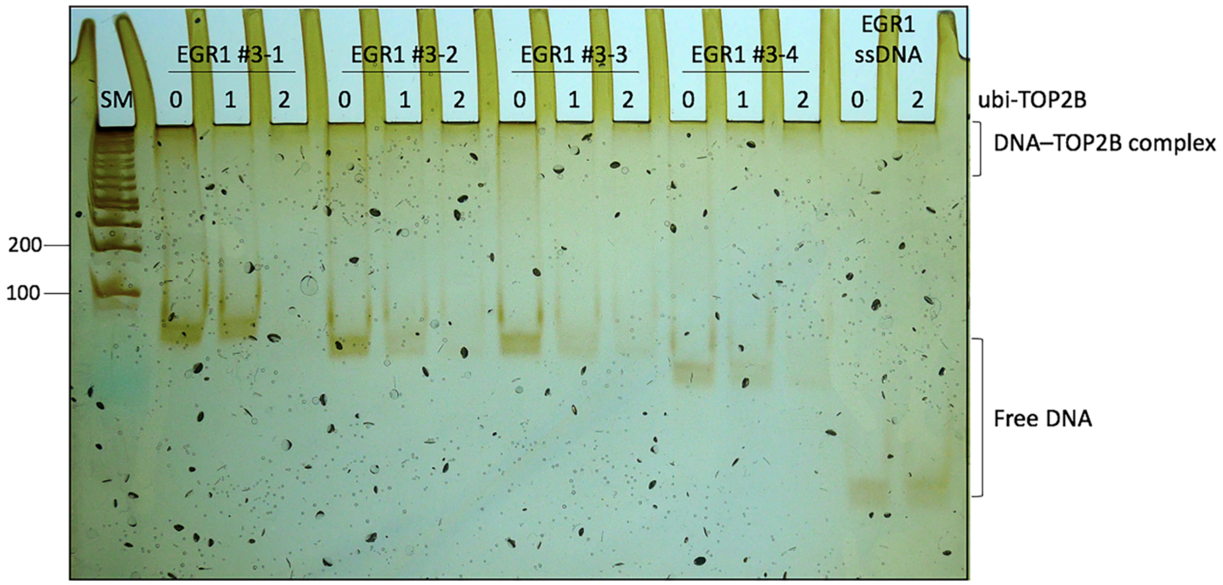
EGR1 #3-1
EGR1 #3-2
EGR1 #3-3
EGR1 #3-4
EGR1
ssDNA
SM
0 1 2 0 1 2 0 1 2 0 1 2 0 2
ubi-TOP2B
DNA–TOP2B complex
200
100
Free DNA

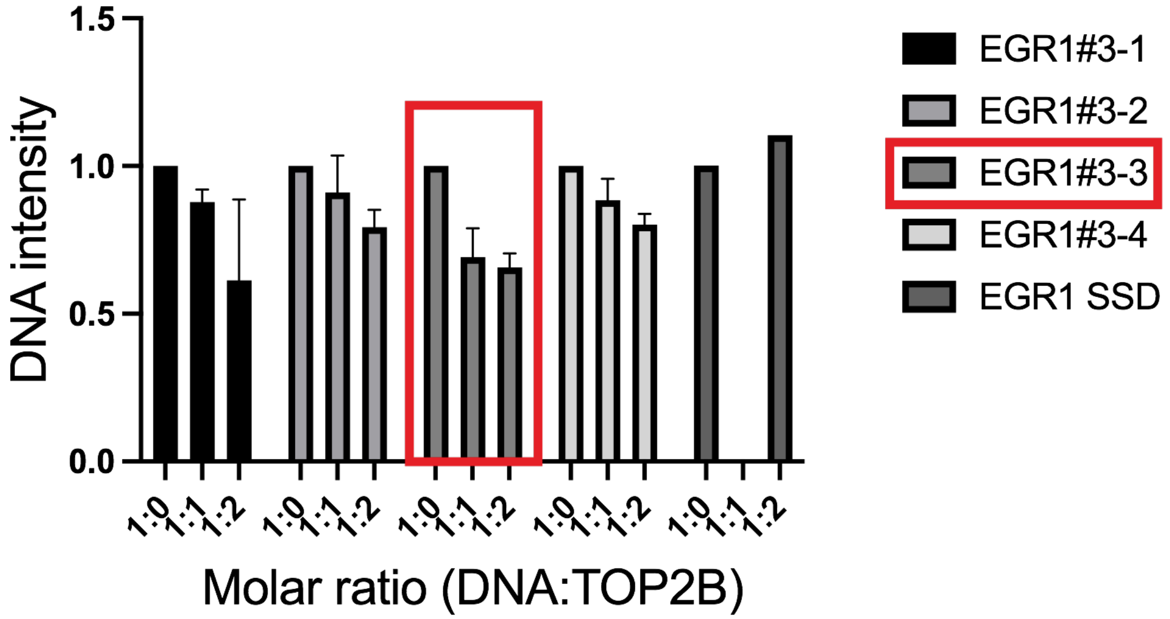
1.5
1.0
0.5
0.0
DNA intensity
1:0 1:1 1:2 1:0 1:1 1:2 1:0 1:1 1:2 1:0 1:1 1:2 1:0 1:1 1:2
Molar ratio (DNA:TOP2B)
EGR1#3-1
EGR1#3-2
EGR1#3-3
EGR1#3-4
EGR1 SSD

**Supplementary Figure 3. Electrophoretic mobility shift assay (EMSA) using TOP2B and *EGR1* TSS fragments.**

(**A**) EMSA results showing the *EGR1* TSS fragment (–132 to +62, EGR1 TSS #3) that was identified to have the strongest affinity to TOP2B from our previous study (Bunch et al. Open Biology 2021). DNA, 100 ng; ubiquitinated TOP2B ($TOP2B^{ubi}$) or deubiquitinated TOP2B ($TOP2B^{deubi}$) titration of 0, 150, 300, 450, and 600 ng. The gel was silver-stained to visualize both DNA and protein.
(**B**) *EGR1* TSS #3 was further dissected into 4 fragments, generating EGR1 #3-1, #3-2, #3-3, and #3-4. The four fragments were compared for binding to TOP2B, using EMSA (Bunch et al. Open Biology 2021). The reactions were separated on the DNA polyacrylamide gel and the resultant gel was stained with silver nitrate. The top panel shows a representative gel and the bottom graph shows the averages and standard deviations from two independent EMSA experiments. Control, *EGR1* TSS ssDNA (or SSD). Per a reaction, 100 ng of DNA was used. 0, no protein; 1 and 2, DNA : TOP2B = 1:1 or 1:2 molar ratio.

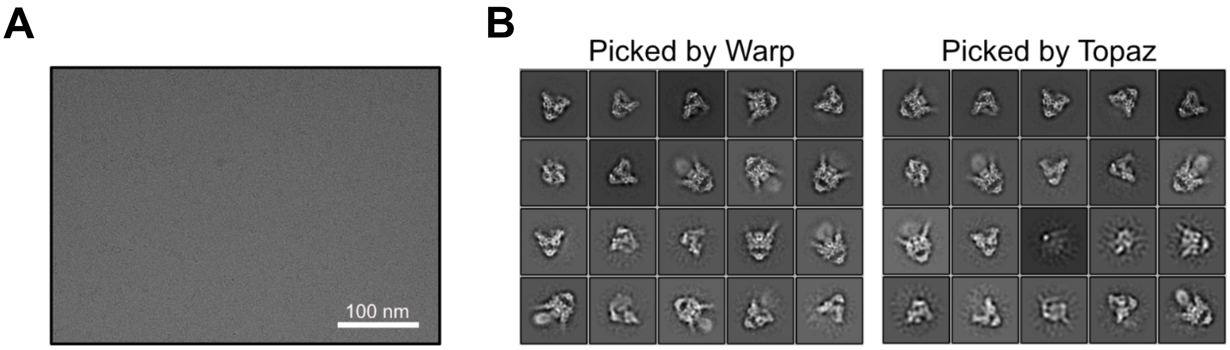


C

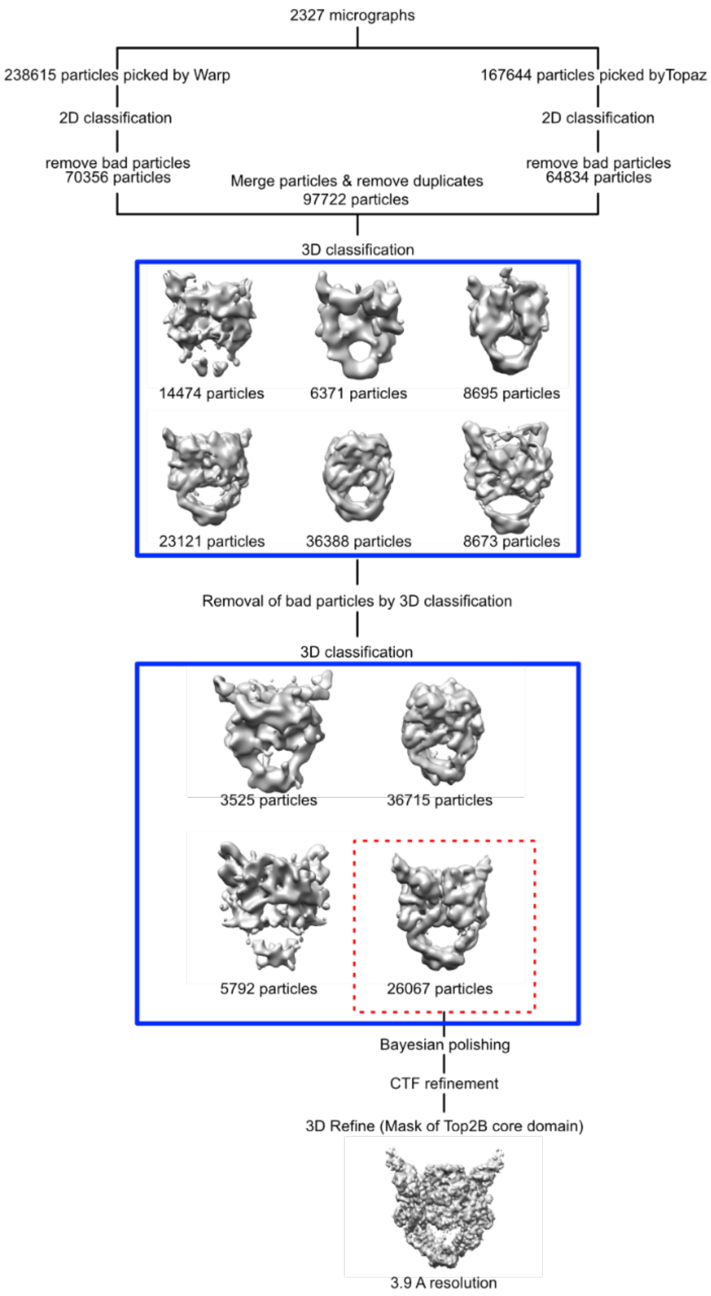
2327 micrographs
238615 particles picked by Warp
167644 particles picked byTopaz
2D classification
2D classification
remove bad particles
70356 particles
remove bad particles
64834 particles
Merge particles & remove duplicates
97722 particles
3D classification
14474 particles
6371 particles
8695 particles
23121 particles
36388 particles
8673 particles
Removal of bad particles by 3D classification
3D classification
3525 particles
36715 particles
5792 particles
26067 particles
Bayesian polishing
CTF refinement
3D Refine (Mask of Top2B core domain)
3.9 A resolution

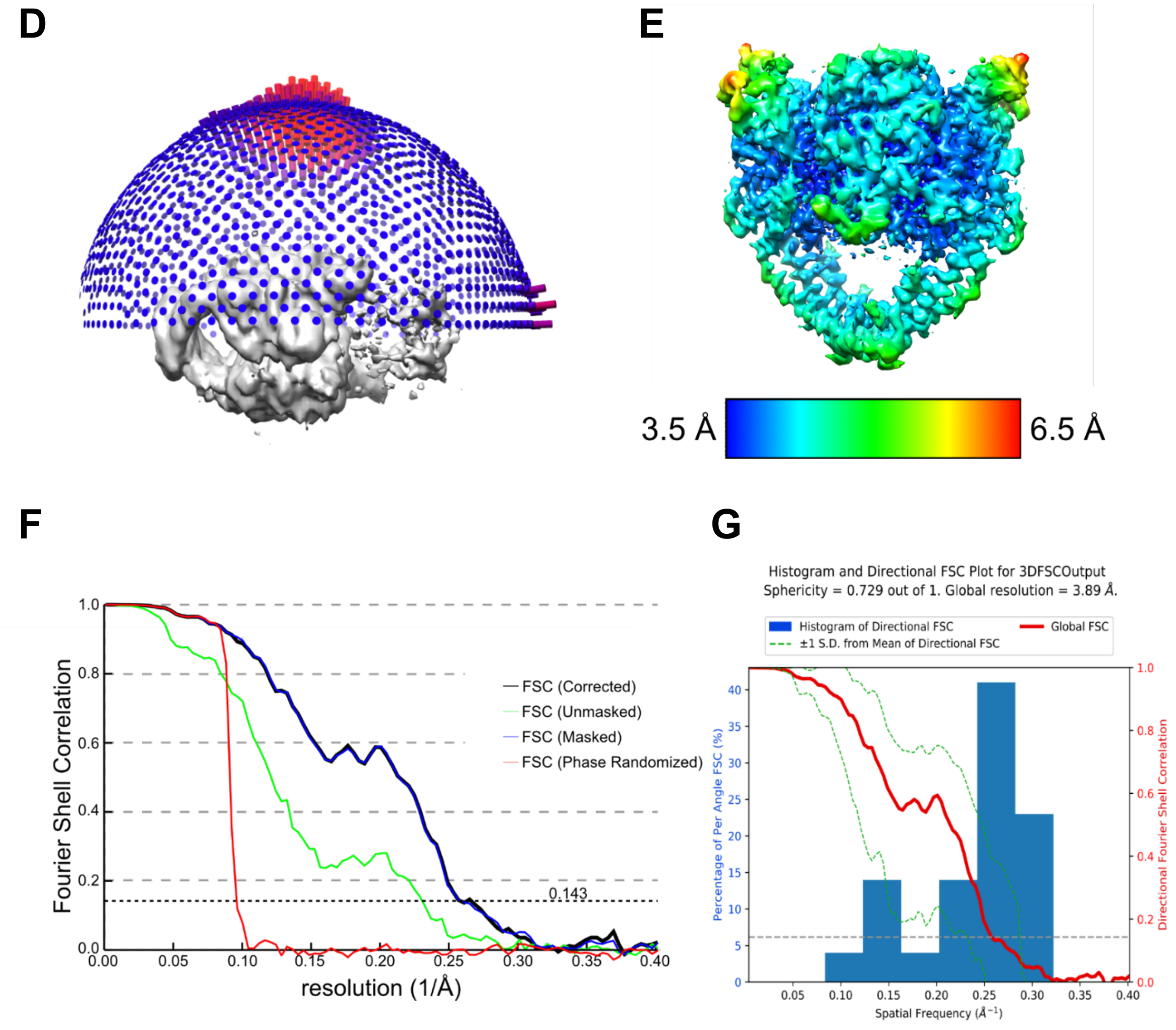


**Supplementary Figure 4. Cryo-EM data collection and image processing.**

(**A**) Representative cryo-EM micrograph.
(**B**) Representative 2D class averages from the reference-free 2D classification calculated after removing bad particles.
(**C**) Flowchart of the image processing.
(**D**) Euler distribution plot of the overall reconstruction.
(**E**) Local resolution estimation of the cryo-EM reconstruction. The map is colored according to the local resolution values. Local resolutions are estimated using the RELION postprocess.
(**F**) Fourier shell correlation curves for the TOP2B-ternary complex. The overall resolution is estimated to be 3.9 Å at an FSC = 0.143.
(**G**) 3DFSC plots of Top2B-DNA-Etoposide complex. The overall resolution is estimated to be 3.9 Å at an FSC = 0.143 with a sphericity of 0.729.

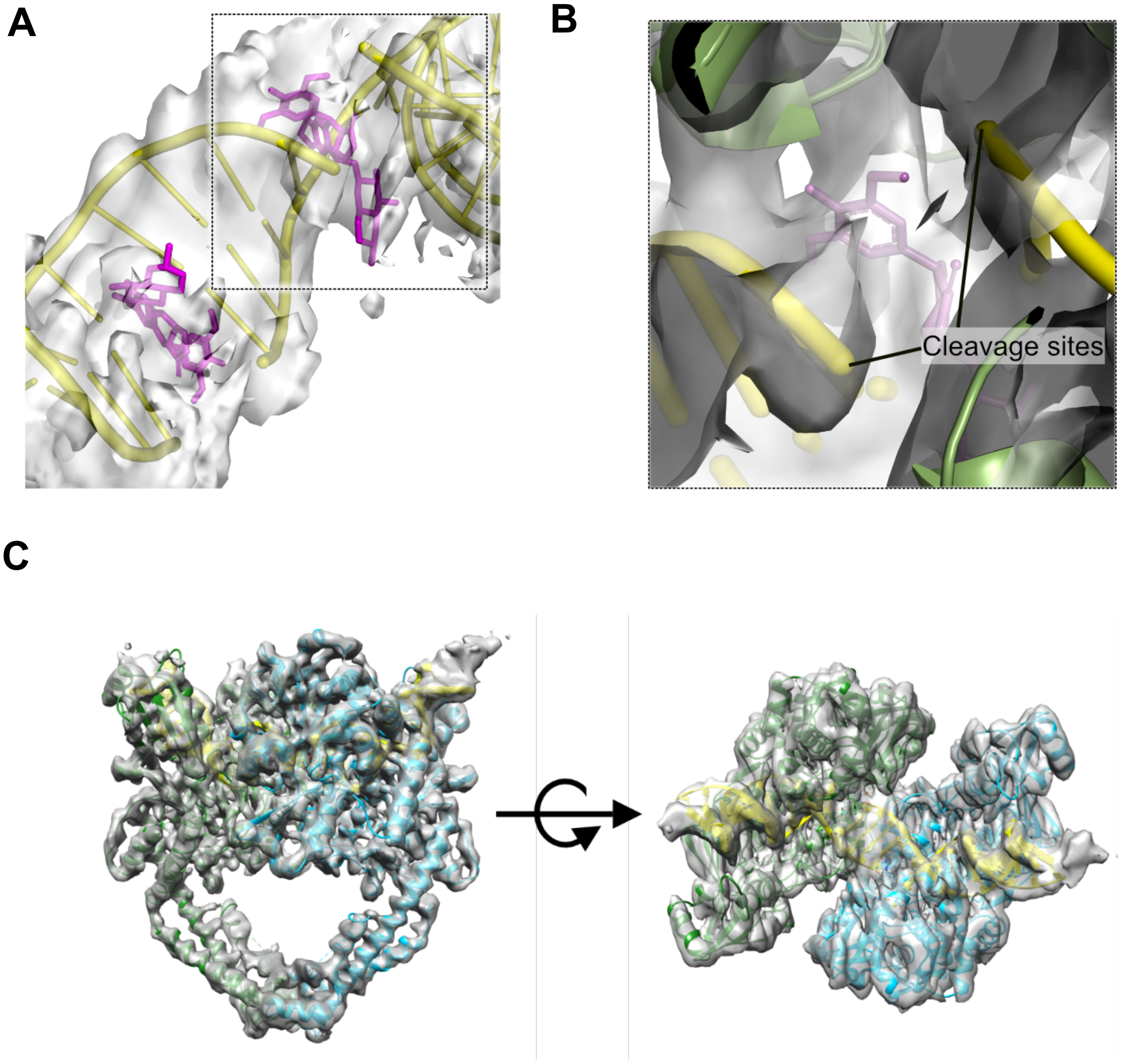
A
B
Cleavage sites
C

**Supplementary Figure 5. Density maps and models of TOP2B-DNA-etoposide complex.**

(**A**) Density map around the etoposide molecules and the cleavage sites of DNA. Etoposide molecules are shown as magenta sticks. DNA molecules are shown as yellow tubes.
(**B)** A close-up view of the square region in the panel A.
(**C**) Density of TOP2B-DNA-etoposide complex without imposing C2 symmetry. The ribbon model of TOP2B-DNA-Etoposide complex refined using the density with imposing C2 symmetry is superposed. There are no notable differences between the densities with and without imposing C2 symmetry.

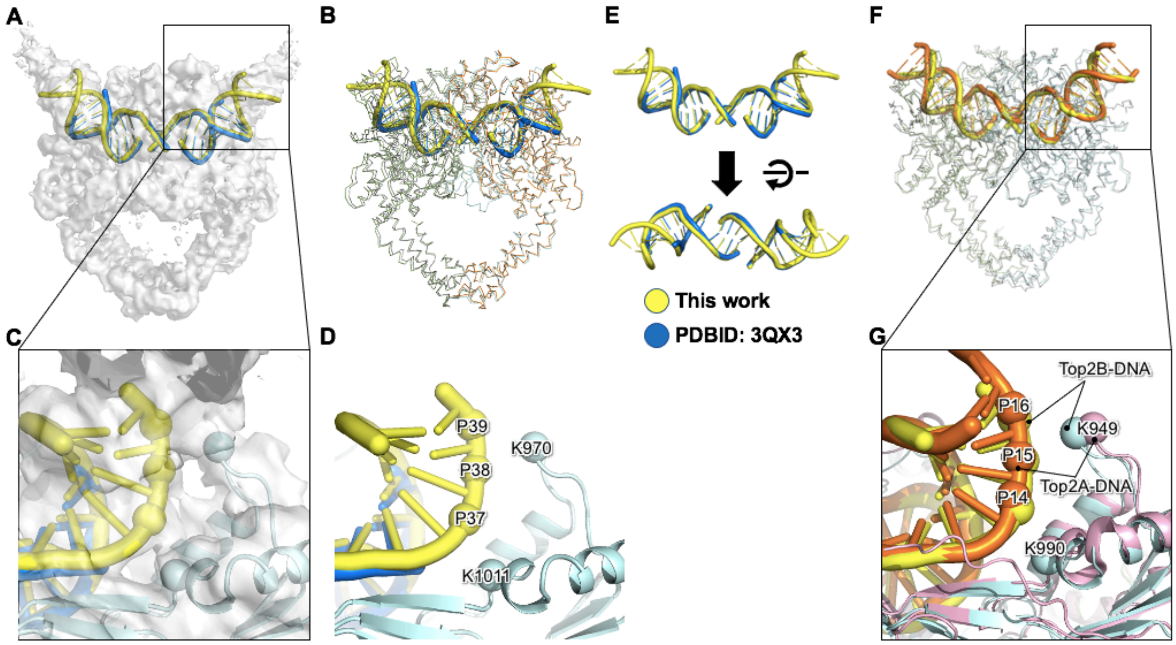


**Supplementary Figure 6. Comparison of other TOP2 structures**.

(**A**) The cryo-EM density map of the TOP2B-DNA-etoposide complex is shown with a cartoon model of DNA colored yellow. The DNA model in the crystal structure of Top2B-DNA-etoposide complex (PDB 3QX3) is overlayed and colored blue.
(**B**) The backbone-trace model of the current TOP2B-DNA-etoposide complex was superimposed with that of the crystal structure (PDB 3QX3) by one of the TOP2B subunits (the right half of the molecules).
(**C**) A close-up view of the region enclosed by a rectangle in panel A. The viewing angle was changed for the visibility. The Top2B protein is represented as a light blue cartoon model. The Cα atoms of Top2B (K970 and K1011) and the DNA phosphate atoms (P37, P38, and P39) are shown as spheres.
(**D**) The cryo-EM density was removed from panel C. The atoms shown as spheres are labeled.
(**E**) Comparison of DNAs in panels A and B.
(**F**) The TOP2B-DNA-etoposide complex structure was superimposed with the TOP2A-DNA-etoposide complex structure by one of the protein subunits. The TOP2 proteins are shown as backbone-trace models. The DNA molecules in the TOP2B and the TOP2A complexes are represented as yellow and orange tubes, respectively.
(**G**) A close-up view of the region enclosed by a rectangle in panel F. The TOP2B and TOP2A proteins are represented as a light blue and pink cartoon models, respectively.

**A**

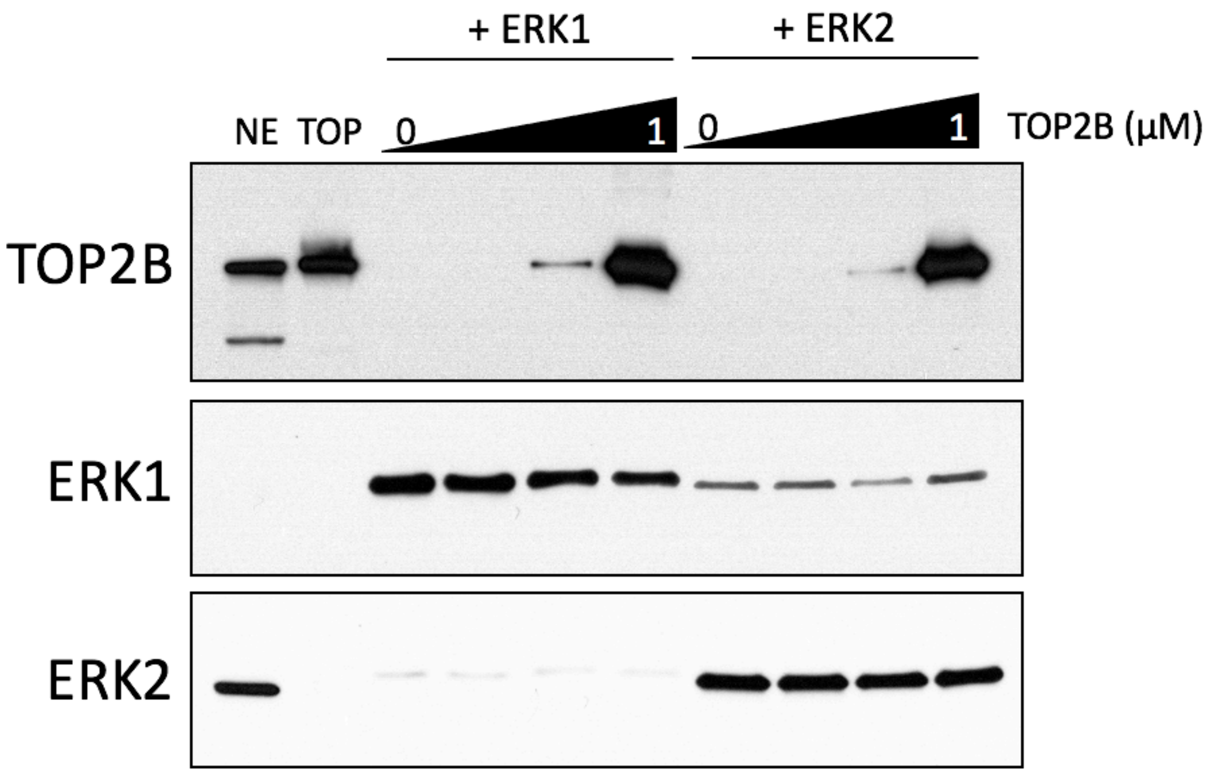


**Supplementary Figure 7. Representative immunoblotting data for DNA relaxation assays**.

(**A**) TOP2B was used at concentrations of 0, 0.005, 0.05, and 1 μM and ERK1 and ERK2 was added at 0.2 μM per each reaction for DNA relaxation assays. One tenth of the reaction was assays through immunoblotting. NE, HeLa nuclear extracts; TOP, TOP2B used for the assay. ERK1 antibody showed some cross-reactivity to ERK2.

### Supplementary Table 1.

The sequences of oligos and primers used in this study

| | |
|---|---|
| **EGR1 template and cloning** | |
| **EGR1 –423 Biotin-Forward** | 5Biotin-GG AGC AGG AAG GAT CCC CCG CCG GAA CAA CCC |
| **EGR1 +332-Reverse** | CCG AAC GGG TCA GAG ATC TGC AGC GGG GAC ATC AGC |
| **EGR1-ELK1$^{mut}$-Forward** | GGA GTG GCC CGA TAT GGC CCG GCC TAA ATT AAC TCT GGG AGG AGG GAA GAA GGC GG |
| **EGR1-ELK1$^{mut}$-Reverse** | CCG CCT TCT TCC CTC CTC CCA GAG TTA ATT TAG GCC GGG CCA TAT CGG GCC ACT CC |
| **ELK1 Cloning-Forward** | CGC CAT ATG ATG GAC CCA TCT GTG ACG CTG TGG CAG TTT CTG CTG C |
| **ELK1 Cloning-Reverse** | CGA TCT CGA GTT AAT GAT GAT GAT GAT GGT GTG GCT TCT GGG GCC CTG GGG AGA GC |
| **ERK2 GA Insertion-Forward** | GCG GCG GCG GCG GGT GCA GGC CCG GAG ATG GTC |
| **ERK2 GA Insertion-Reverse** | GAC CAT CTC CGG GCC TGC ACC CGC CGC CGC CGC |
| **ERK2 L46V QC-Forward** | GGT TTG TTC TGC TTA TGA TAA TGT CAA CAA AGT TCG AGT TGC |
| **ERK2 L46V QC-Reverse** | GCA ACT CGA ACT TTG TTG ACA TTA TCA TAA GCA GAA CAA ACC |
| **qRT-PCR** | |
| **EGR1-Forward** | CTT CAA CCC TCA GGC GGA CA |
| **EGR1-Reverse** | GGA AAA GCG GCC AGT ATA GGT |
| **FOS-Forward** | CAC AGA CCC AGG CCT GGC TCA ACA TGC TAC |
| **FOS-Reverse** | CAC CAG GCT GTG GGC CTC AAG GAC TTG AAA GC |
| **HSP70-Forward** | ATG TCG GTG GTG GGC ATA GA |
| **HSP70-Reverse** | CAC AGC GAC GTA GCA GCT CT |
| **ACTIN-Forward** | GCC GAC AGG ATG CAG AAG GAG ATC A |
| **ACTIN-Reverse** | AAG CAT TTG CGG TGG ACG ATG GA |
| **EMSA & CryoEM** | |
| **EGR1 –133 to –71 (EGR1 #3-1) duplex 1** | GTC GTG ACG TAC ATG GCC ATA TAT GGG AAG CAG GAA GCC CTA ATA TGG CAG GAC CGG CCG GGA |
| **EGR1 –133 to –71 (EGR1 #3-1) duplex 2** | TCC CGG CCG GTC CTG CCA TAT TAG GGC TTC CTG CTT CCC ATA TAT GGC CAT GTA CGT CAC GAC |
| **EGR1 –70 to –21 (EGR1 #3-2) duplex 1** | CCC GGA TCC GCC TCT ATT TGA AGG GTC TGG AAC GGC ACG GGT CCG CCT CC |
| **EGR1 –70 to –21 (EGR1 #3-2) duplex 2** | GGA GGC GGA CCC GTG CCG TTC CAG ACC CTT CAA ATA GAG GCG GAT CCG GG |
| **EGR1 –20 to +30 (EGR1 #3-3) duplex 1** | ATG GCG GCG GCG GCT CCC CAA GTT CTG CGC GCT GGG ATC TCT CGC GAC TC |
| **EGR1 –20 to +30 (EGR1 #3-3) duplex 2** | GAG TCG CGA GAG ATC CCA GCG CGC AGA ACT TGG GGA GCC GCC GCC GCC AT |

| | |
|---|---|
| **EGR1 +31 to +70 (EGR1 #3-4) duplex 1** | GGG CCG GTC CTG CGG CGG CGG AAG CTG GCT GCG GCG GCG G |
| **EGR1 +31 to +70 (EGR1 #3-4) duplex 2** | CCG CCG CCG CAG CCA GCT TCC GCC GCC GCA GGA CCG GCC C |
| **EGR1 TSS ssDNA** | CCT GCT TCC CAT ATA TGG CCA TGT ACG |
| **ChIP-qPCR** | |
| **EGR1 TSS-Forward** | CCT GCT TCC CAT ATA TGG CCA TGT ACG |
| **EGR1 TSS-Reverse** | CCT GCG GCG GCG GAA GCT GGC TGC |
| **FOS TSS-Forward** | GAC CGT GCT CCT ACC CAG CTC TGC TCC ACA GCG CCC |
| **FOS TSS-Reverse** | GAG TGG TAG TAA GAG AGG CTA TCC CCG GCC |
| ***In vitro* transcription** | |
| **EGR1 +1 Forward** | GCG CAG AAC TTG GGG AGC CGC CGC CGC C |
| **EGR1 +332-Reverse** | CCG AAC GGG TCA GAG ATC TGC AGC GGG GAC ATC AGC |

### Supplementary Table 2.

### Cryo-EM information

| | |
|---|---|
| **Data collection and processing** | |
| Magnification | 105000 |
| Voltage (kV) | 300 |
| Electron exposure (e−/A2) | 49 |
| Defocus range (um) | 1.2-2.5 |
| Pixel size (Å) | 0.83 |
| Symmetry imposed | C2 |
| Initial particle images (no.) | 238615 |
| Final particle images (no.) | 26067 |
| Map resolution (Å) | 3.9 (FSC = 0.143), 4.7 (FSC = 0.5) |
| **Refinement** | |
| Initial model used | alphafold2 (UniProt Q02880) |
| Map sharpening B factor ($Å^2$) | -102.69 |
| **Model composition** | |
| Non-hydrogen atom | 13036 |
| Protein residues | 1460 |
| DNA residues | 56 |
| Ligands | 2 EVP, 2 MG |
| **B factors ($Å^2$)** | |
| Protein residues | 164.16 |
| DNA residues | 181.07 |
| Ligands | 169.93 |
| **R.m.s. deviations** | |
| Bond length (Å) | 0.003 |
| Bond angles (°) | 0.576 |
| **Validation** | |
| MolProbity score | 1.87 |
| Clashscore | 13.05 |
| Poor rotames (%) | 0 |
| **Ramachandran plot** | |
| Favored (%) | 96.35 |
| Allowed (%) | 3.65 |
| Disallowed (%) | 0 |
